# Bracketing phenotypic limits of mammalian hybridization

**DOI:** 10.1101/310789

**Authors:** Yoland Savriama, Mia Valtonen, Juhana Kammonen, Pasi Rastas, Olli-Pekka Smolander, Annina Lyyski, Teemu J. Häkkinen, Ian J. Corfe, Sylvain Gerber, Isaac Salazar-Ciudad, Lars Paulin, Liisa Holm, Ari Löytynoja, Petri Auvinen, Jukka Jernvall

**Affiliations:** Developmental Biology Program, Institute of Biotechnology, University of Helsinki, P.O. Box 56, FIN-00014 Helsinki, Finland.; Department of Environmental and Biological Sciences, University of Eastern Finland, P.O. Box 111, FIN-80101 Joensuu, Finland.; Genome Biology Program, Institute of Biotechnology, University of Helsinki, P.O. Box 56, FIN-00014 Helsinki, Finland.; Institut Systématique Evolution Biodiversité (ISYEB), Muséum national d’Histoire naturelle, CNRS, Sorbonne Université, EPHE, 45 rue Buffon, CP 50, 75005 Paris, France.; Departament de Genètica i Microbiologia, Universitat Autònoma de Barcelona, 08193 Cerdanyola del Vallès, Spain.; Faculty of Biological and Environmental Sciences, University of Helsinki, P.O. Box 56, FIN-00014 Helsinki, Finland.

## Abstract

An increasing number of mammalian species have been shown to have a history of hybridization and introgression based on genetic analyses. Only relatively few fossils, however, preserve genetic material and morphology must be used to identify the species and determine whether morphologically intermediate fossils could represent hybrids. Because dental and cranial fossils are typically the key body parts studied in mammalian paleontology, here we bracket the potential for phenotypically extreme hybridizations by examining uniquely preserved cranio-dental material of a captive hybrid between gray and ringed seals. We analyzed how distinct these species are genetically and morphologically, how easy it is to identify the hybrids using morphology, and whether comparable hybridizations happen in the wild. We show that the genetic distance between these species is more than twice the modern human-Neanderthal distance, but still within that of morphologically similar species-pairs known to hybridize. In contrast, morphological and developmental analyses show gray and ringed seals to be highly disparate, and that the hybrid is a predictable intermediate. Genetic analyses of the parent populations reveal introgression in the wild, suggesting that gray-ringed seal hybridization is not limited to captivity. Taken together, gray and ringed seals appear to be in an adaptive radiation phase of evolution, showing large morphological differences relative to their comparatively modest genetic distance. Because morphological similarity does not always correlate with genetic distance in nature, we postulate that there is considerable potential for mammalian hybridization between phenotypically disparate taxa.

## Introduction

Although hybridization has been extensively examined in the context of speciation, the role of interbreeding leading to introgression and admixture of phenotypic traits is attracting increasing attention (Ackermann et al., 2006; Mallet, 2007; Abbott et al., 2013; Polly et al., 2013; Pennisi, 2016; Árnason et al., 2018; Lamichhaney et al., 2018). In studies of human evolution, genetic evidence has implicated introgression between different lineages of *Homo* (Green et al., 2010; Meyer et al., 2012), and an increasing number of mammalian fossils are suggested to retain signs of interbreeding by palaeogenomic studies (Enk et al., 2011; Fu et al., 2011; Wecek et al., 2017). Hybridization can complicate the assignment of fossil specimens to specific taxa (Trinkaus et al., 2003; Trinkaus, 2007; Polly et al., 2013; Martinón-Torres et al., 2017), and currently it remains unclear how phenotypically different or disparate taxa might be expected to hybridize. Even if hybrids between morphologically dissimilar species have reduced fertility, they may still be preserved in the fossil record, especially if hybridization is fairly common as in active hybrid zones (e.g., Mallet, 2007; Polly et al., 2013; Shurtliff, 2013; Pallares et al., 2016). To assess the maximum morphological range of potential hybridization in the fossil record, analyses of hybrids between morphologically disparate taxa are required. One such hybridization, with uniquely preserved cranial material, has been reported to have occurred in Stockholm zoo in 1929 between two mammalian species belonging to different genera; the gray seal (*Halichoerus grypus*) and the ringed seal (*Pusa hispida*) (Lönnberg, 1929). This incident raises questions as to how distinct these species are genetically, how distinct these species are phenotypically, how easy it is to identify the hybrids using morphology, and whether comparable hybridizations happen in the wild. Addressing all these questions together allows one to estimate the ‘hybrid bracket’, or the overall potential for hybridization in mammalian evolution.

## Results

### Validating the seal hybrid and the genetic context of the hybridization

The seal born in 1929 was immediately concluded to be a hybrid because at the time the seal pond housed only three adult seals; two male gray seals and one female ringed seal (Figure 1). The newborn was found dead and although no malformations were reported (Lönnberg, 1929), both biological and husbandry related causes of death remain a possibility. First, to verify the hybridization and validate the identity of the museum specimen, we sequenced its genome (see Materials and Methods, Table supp. 1) together with the genomes of its parental species, Baltic gray seals (*n* = 10) and Baltic ringed seals (*n* = 9) (see Materials and Methods). A genetic admixture analysis shows that the hybrid shares roughly 50% of its genome with both species, confirming that it is indeed a hybrid between the gray and the ringed seal (Figure 1C).

**Figure 1.**
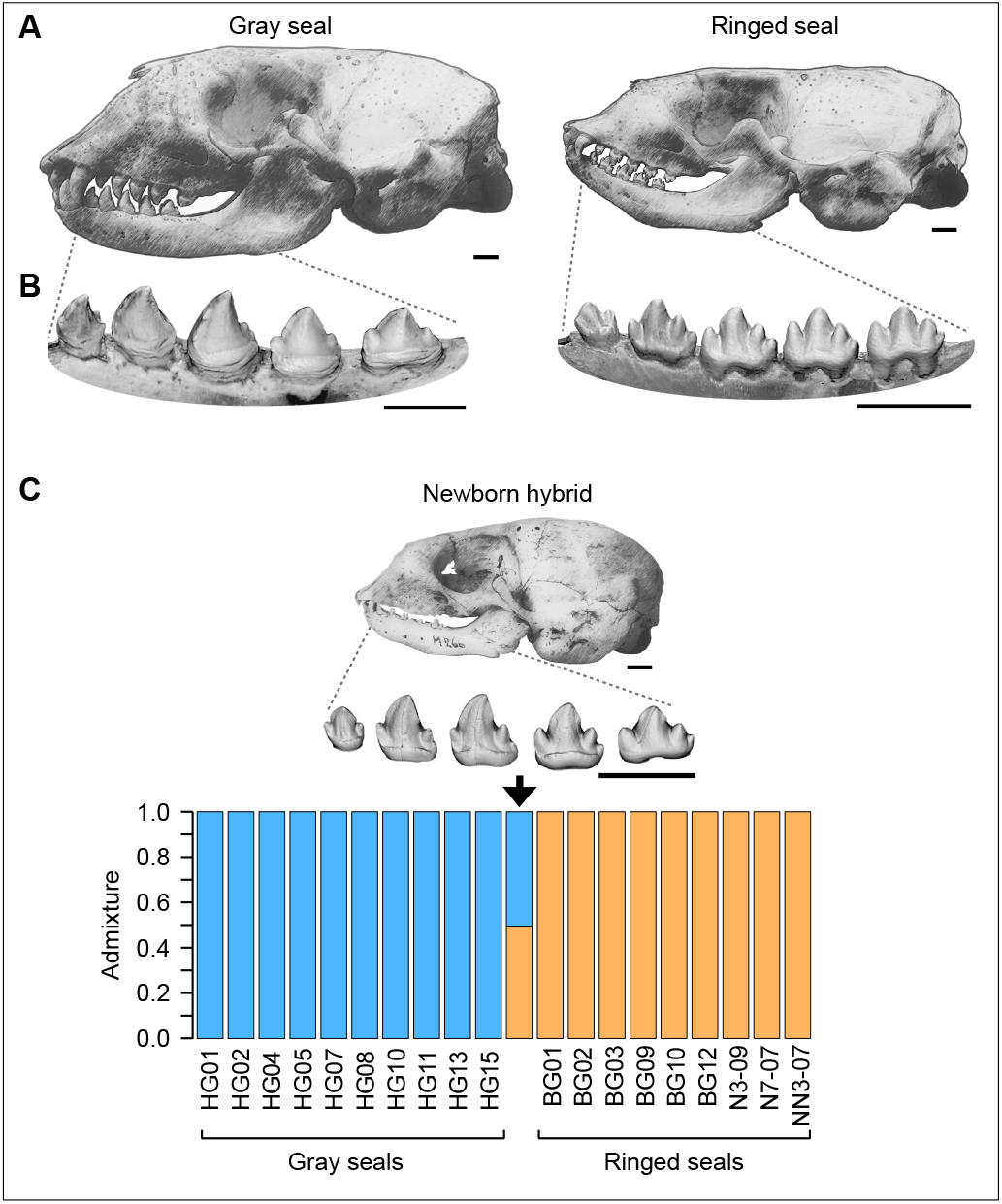
Gray and ringed seal cranio-dental morphologies show at least genus-level differences and genetic data validates the hybrid specimen. **(A)** The gray seal skull has a prominent snout with a straight profile whereas the overall cranial morphology of ringed seals is relatively gracile. **(B)** The gray seal lower postcanines have robust, fang-shaped central cusps with small and variably present accessory cusps. There is a marked gradation of tooth shapes along the jaw, and gray seal teeth are approximately 40% larger (anteroposteriorly) than ringed seal teeth. The ringed seal lower postcanines have typically four slender cusps on the four large postcanines of the lower jaw. Upper teeth show comparable but less disparate differences in cusp number. Gray seals have pronounced sexual dimorphism and the skull illustrations are of females. Postcanine dentitions, showing obliquely lingual views, have no marked sexual dimorphism. **(C)** The hybrid specimen is a newborn skull and mandible with erupting, but fully formed teeth. Genetic analysis from hybrid dental pulp shows close to 50% admixture between gray (50.51%) and ringed (49.49%) seals. Analysis shown (K=2) was performed using NGSadmix of ANGSD with 3.6 million markers (see Materials and Methods). Scale bars, 10 mm.

The exact phylogenetic position and distinctiveness of the gray seal in relation to the ringed seal and other related taxa has been problematic (Fulton and Strobeck, 2010; Nyakatura and Bininda-Emonds, 2012; Berta et al., 2018), leaving open the question how genetically distinct the species really are. To approximate the genetic context of the seal hybridization, we computed a genome-wide estimate of neutral genetic distance between gray and ringed seals (see Materials and Methods), and contrasted this with species pairs well known to hybridize; lion-tiger (*Panthera leo–P. tigris*) and domestic donkey–horse (*Equus asinus–E. ferus*) (see Materials and Methods, Table supp. 2). Lion and tigers readily hybridize in captivity (Vella et al., 1999), and as both they and seals are members of Carnivora, provide an appropriate comparison. Whereas these carnivoran species-contrasts have the same number of chromosomes [2n = 32 in seals (Corfman and Richart, 1964; Árnason, 1970) and 2n = 38 in felids (Vella et al., 1999)], donkey and horse differ in their chromosome number and their hybrids, known as mules and hinnies, are generally infertile [2n = 62 in donkeys and 2n = 64 in horses (Jónsson et al., 2014)]. To place the results in the context of *Homo* lineage, human and Neanderthal were also included in the analysis, as were some additional outgroups (Table supp. 2).

To obtain a robust proxy of neutral sequence evolution, we used the fourfold degenerate sites of 4,045 protein-coding genes, present as single copy in each species (see Materials and Methods). Analyses of 623,391 fourfold degenerate sites from genes orthologous for all the eight species show that the gray-ringed seal genetic distance is roughly 49% and 26% of the lion-tiger and donkey-horse distances, respectively (Figure 2, Table 1). These distances appear robust when larger or smaller sets of species are compared (Table supps 3, 4). The comparatively short genetic distance between the gray and ringed seals is noteworthy because unlike in the hybridizing lion-tiger and donkey-horse comparisons, the dental and cranial morphologies of the two seal species have historically warranted a genus level distinction (Fulton and Strobeck, 2010; Nyakatura and Bininda-Emonds, 2012; Berta et al., 2018) in contrast to the species level distinctions of the lion-tiger and donkey-horse pairs.

**Figure 2.**
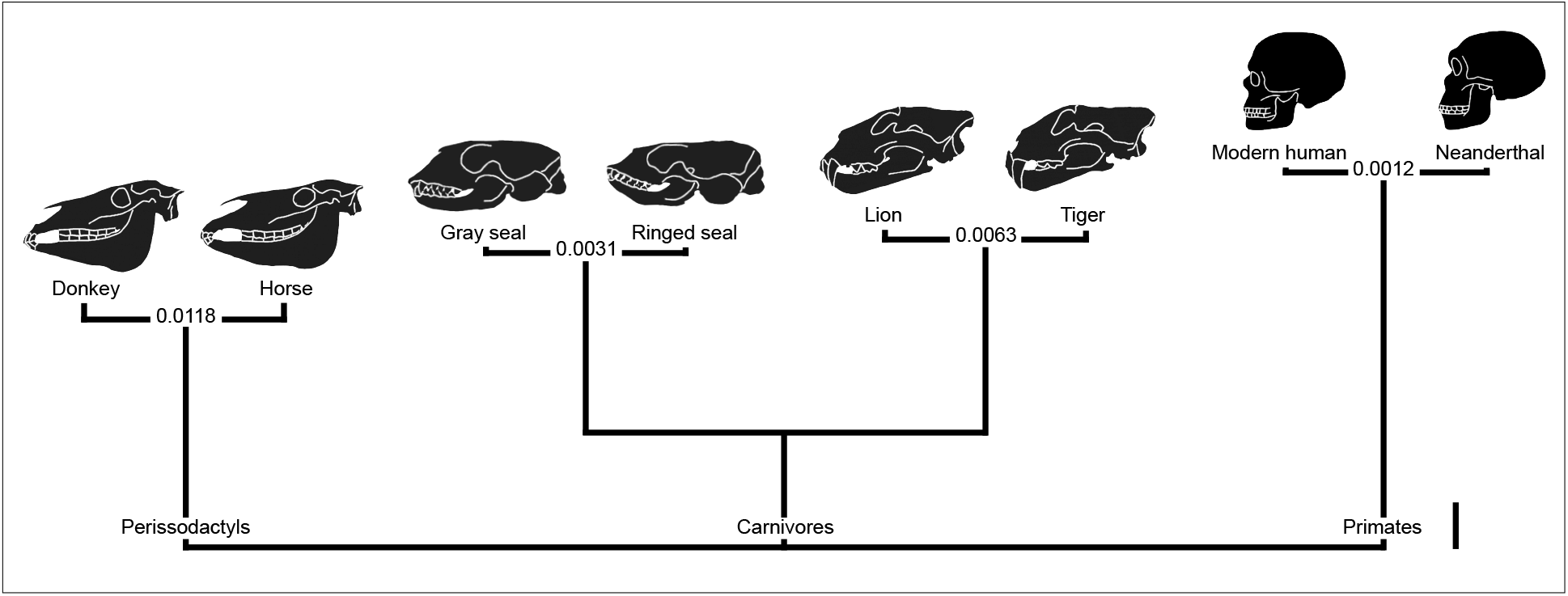
Bracketing the genetic distance of gray and ringed seals with other hybridizing taxa. Lion-tiger and donkey-horse distances (substitutions per fourfold degenerate site) are longer, and human-Neanderthal distance is shorter than the gray-ringed seal distance. Of the hybridizing species pairs, only gray and ringed seals are in different genera, and have pronounced differences in both dental and cranial morphologies. Pairwise distances (T92 model) and a phylogenetic tree were built using fourfold degenerate sites of 4,045 orthologous genes. For details and all the distances, see Materials and Methods and Table 1. Scale bar, 0.02 substitutions per ffd. The crania shown are not to scale. The horse cranium is of *E. f. przewalskii*.

**Table 1.** Genetic distances between hybridizing species (substitutions per fourfold degenerate site).

|  | Human | Neanderthal | Tiger | Lion | Horse | Donkey | Ringed seal |
| --- | --- | --- | --- | --- | --- | --- | --- |
| Neanderthal | 0.0012 |  |  |  |  |  |  |
| Tiger | 0.2596 | 0.2596 |  |  |  |  |  |
| Lion | 0.2599 | 0.2598 | 0.0063 |  |  |  |  |
| Horse | 0.2369 | 0.2368 | 0.2164 | 0.2165 |  |  |  |
| Donkey | 0.2366 | 0.2365 | 0.2159 | 0.2160 | 0.0118 |  |  |
| Ringed seal | 0.2539 | 0.2539 | 0.1541 | 0.1542 | 0.2084 | 0.2079 |  |
| Gray seal | 0.2540 | 0.2539 | 0.1544 | 0.1544 | 0.2086 | 0.2082 | 0.0031 |

Based on 1,526,260 codons, 623,391 fourfold degenerate sites, and 4,045 orthologous genes.

Even though the gray-ringed seal genetic distance is shorter than the carnivore and perissodactyl contrasts (Figure 2), it is still considerably longer than the hominin contrast. Compared to the modern human-Neanderthal distance, the gray-ringed seal distance is roughly two and a half times greater (258%, Table 1), suggesting that many fossil hominins are within the genetic hybrid bracket. Since several of the potentially hybridizing hominins are also phenotypically different, next we analyzed the dental and cranial distinctiveness of the two seal species and their hybrid. These phenotypic structures are the key features in many paleontological studies.

### Analyzing the teeth of the hybrid

Mammalian tooth shape is fully formed prior to function with no remodeling other than wear after mineralization, thereby providing relatively direct information about development. In addition, seals have vestigial deciduous dentitions and are born with an erupting permanent dentition. Even though the newborn hybrid was found dead, the precocious state of seal dental development allows us to compare the hybrid with adult seals (Figure 1C). The original description of the hybrid by Lönnberg (1929), while detailed, was not quantitative and therefore we 3D-reconstructed the hybrid dentition from microCT scans (see Materials and Methods, Figure 3A, Figure supp. 1). Here we focus on the lower postcanine dentitions as they show the largest range of variation and have been studied previously (Jernvall, 2000; Salazar-Ciudad and Jernvall, 2010). Whereas seal dentitions are relatively derived from the basic carnivoran pattern, seal postcanine morphologies are reminiscent of various pretribosphenic patterns in mammalian evolution, classified to different families and orders (Luo, 2007; Grossnickle and Polly, 2013).

**Figure 3.**
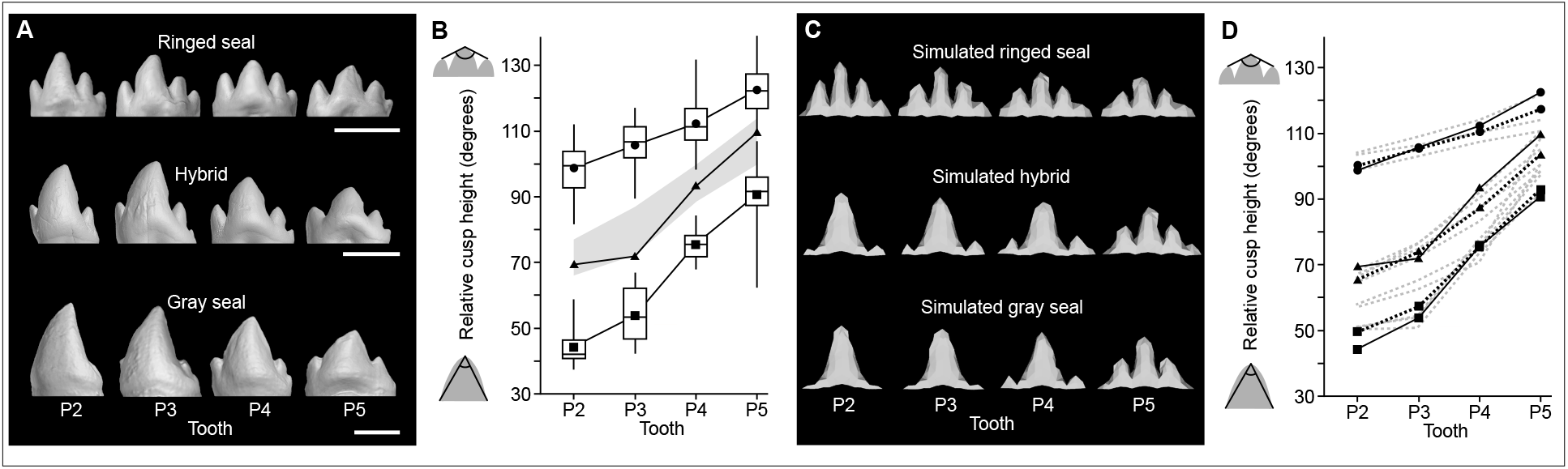
Teeth of the hybrid are morphologically and developmentally intermediate. **(A)** The hybrid has the large central cusps of the gray seal together with relatively prominent accessory cusps reminiscent of the ringed seals. **(B)** The top-cusp angle measure shows that the hybrid teeth (triangles) are intermediate between the ringed (circles, *n* = 43, 50, 49, and 49 for P2, P3, P4, and P5) and gray (squares, *n* = 41, 44, 39, and 53 for P2, P3, P4, and P5) seals with no overlap in P2 to P4, and marginal overlap with ringed seal teeth in P5. Gray area marks 80% of the angles averaged between each gray-ringed seal pair (see Figure supp. 2). Boxes enclose 50% of observations; the median and mean are indicated with a horizontal bar and circle or square, respectively, and whiskers denote range. **(C)** Simulated ringed and gray seal tooth rows that were manually matched with mean real shapes. The tooth row of the simulated hybrid was obtained by averaging the values of the parents. **(D)** The top-cusp angle measure shows that the average-value-simulated hybrid teeth (triangles, black dashed line) are intermediate between the simulated gray (squares, black dashed line) and ringed (circles, black dashed line) seal teeth, and comparable to the real hybrid teeth (triangles, black solid line). Gray dashed lines show hybrid simulation in which maternal or paternal values were tested for each parameter (Figure supp. 3, Table supp. 8). Teeth have been mirrored if needed to show left buccal views. Scale bars, 5 mm.

The overall morphologies of the gray and ringed seal postcanine teeth are markedly different. Ringed seal teeth have three to five slender cusps, and the teeth are generally more similar along the jaw (Figure 1B, Figure 3A, Table supp. 5). In contrast, especially the anterior postcanine teeth of the gray seal have large, fang-shaped central cusps with small, variably present accessory cusps (Figure 1B, Figure 3A, Table supp. 5). These morphological differences reflect the use of smaller fish and invertebrate foods by ringed seals compared to gray seals that, in addition to fish are known to prey on mammals (Stringell et al., 2015; van Neer et al., 2015). Because standard morphometric methods cannot be effectively used to compare teeth with different numbers of tooth cusps, we quantified the shapes by using a top-cusp angle, a measure of relative cusp height used previously in the analyses of seal dentitions (see Materials and Methods) (Jernvall, 2000; Salazar-Ciudad and Jernvall, 2010).

The results show that the hybrid teeth are intermediate between a sample (*n* = 130) of the parent species by having relatively large central cusps, but also relatively prominent accessory cusps (Figure 3A, Figure supp. 2). The distinct and intermediate morphology of the hybrid is most readily visible in the anterior postcanines (P2 and P3) in which the species differences are also the greatest due to the stronger anteroposterior gradation in the gray seal dentition (Figure 3A, B, Figure supp. 2). In addition to shape, tooth size also appears intermediate between the gray and ringed seal (Figure 3A). Because of the intermediate cusp morphologies of the hybrid, next we examined what kind of developmental changes might drive the observed patterns.

### Developmental basis of the hybrid teeth

Genetic regulation of mammalian tooth development is highly conserved (Jernvall and Thesleff, 2012), and reiterative activation of signaling centers, called secondary enamel knots, appear to direct cusp development in all studied mammals (Jernvall and Thesleff, 2012). Consequently, computational modeling of genetic interactions and tissue biomechanics, based on empirical data on mouse tooth development, has been used to model embryological development of the teeth of rodent species (Harjunmaa et al., 2014; Renvoisé et al., 2017), as well as seal teeth (Salazar-Ciudad and Jernvall, 2010). Since neither seal tooth development nor hybridization is amenable to experimentation, we examined whether the hybrid morphology could be produced by modeling development.

First, we generated virtual ringed and gray seal tooth rows (using real population mean shapes, see Materials and Methods) using the ToothMaker-software (Harjunmaa et al., 2014; Renvoisé et al., 2017) (see Materials and Methods). Setting three model parameters to different values was sufficient to model the differences between gray and ringed seal teeth; inhibitor (*Inh*), epithelial growth rate (*Egr*), and anterior bias (*Abi*). These parameters, by regulating the dynamics of development, affect the spacing of cusps, the pointedness of cusps, and the anterior-posterior symmetry of teeth, respectively (Figure supp. 3A, see Materials and Methods). Changes along the tooth row were produced by a constant change in *Egr* in ringed seal models and a constant change in *Egr* and *Inh* in gray seal models (Table supp. 6, see Materials and Methods). The constant parameter changes not only parsimoniously account for well recognized gradual shape changes along the tooth row (Butler, 1939; Van Valen, 1962; Bateson, 1984), but also agree with the inhibitory cascade model that predicts a constant change in tooth size proportions along the jaw (Kavanagh et al., 2007).

To mimic the hybridization between gray and ringed seals, the development of each tooth along the jaw was simulated after adjusting each of the three parameters separating the two species. For each parameter we used either the average parameter value (assuming no dominance) or value adjusted 10% towards each of the parents. Additionally, we simulated teeth by keeping one or two of the parameters at the parent values. We then simulated tooth development using all these parameter value combinations (see Materials and Methods, Figure supp. 3B, Table supps 6–8).

The resulting simulated tooth shapes show that the hybrid cusp patterns can be produced by averaging the three parameters between the modeled gray and ringed seals (Figure 3C). Furthermore, the top-cusp angles of this simulated hybrid tooth row fall between the parent shapes similarly to the real hybrid (Figure 3D, Figure supp. 3B, Table supps 7, 8), suggesting that the regulatory principles of tooth shape are largely conserved between the two seal species.

A key parameter differentiating the gray and ringed seal teeth that is also required to be closest to the average between the parents is the inhibitor (*Inh*), as otherwise the simulated hybrid would have the top-cusp angle close to one of the parents (Figure 3D, Table supps 7, 8, Figure supp. 3B). Experiments on developing mouse teeth have identified sonic hedgehog (SHH) as one candidate molecule for inhibition of new enamel knots and cusps (Harjunmaa et al., 2014). However, because cusp spacing can be altered by tinkering with the activator-inhibitor balance of enamel knots through several molecules, these results do not necessarily implicate a single gene underlying each parameter. Rather, regardless of whether tooth shape is driven by a large number of loci or relatively few loci with major phenotypic effects (for discussion, see Pallares et al., 2016), our simulation results are strongly suggestive that the hybrid dentition is both phenotypically and developmentally a predictable intermediate between the species.

### Analyzing the cranium of the hybrid

Unlike the dentition, the cranium of the newborn hybrid does not allow direct comparison to adult seals. However, differences in cranium morphology can be compared using geometric morphometrics across species and age groups (Klingenberg, 2010), and we digitized 46 3D-landmarks from newborn to adult gray and ringed seals (Figure supp. 4, Table supp. 9, *n* = 116, Movie supp. 1, see Materials and Methods) to depict overall changes in skull shape.

The results show that gray and ringed seal crania have distinct developmental trajectories, and the newborn crania from each species are already well separated in the morphospace with no overlap (*p* < 0.0001, permutation test on Procrustes distance with 10,000 rounds, Figure 4). The hybrid cranium is positioned between the two species, and in the proximity of the geometric morphometric mean of newborn gray and ringed seals (Figure 4). Therefore, as is the case for the dentition, the overall morphology of the cranium appears to be an intermediate between the species, a result agreeing with recent results on hybrids between different mouse strains (Warren et al., 2018).

**Figure 4.**
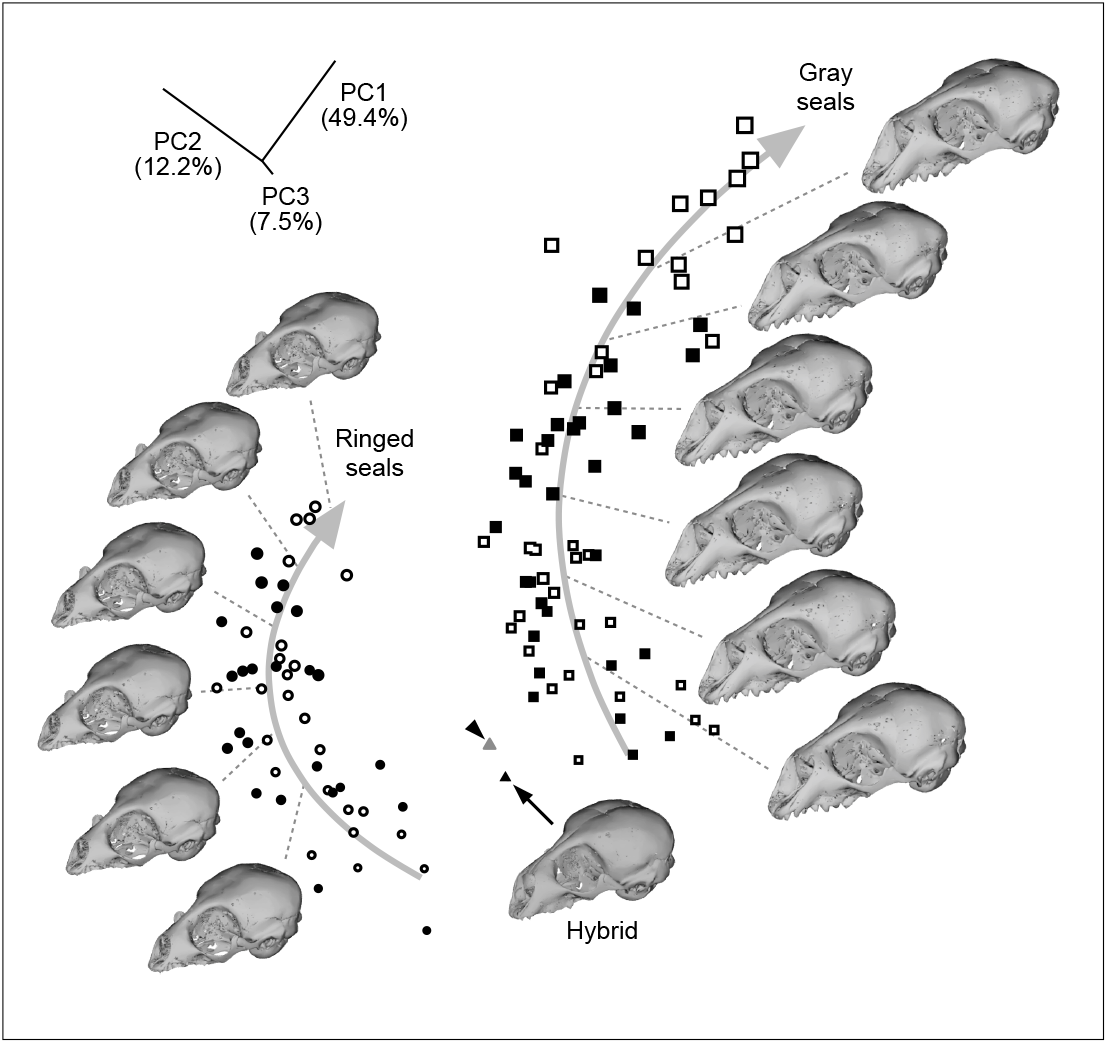
Ringed and gray seal crania are distinct at the birth and the hybrid is intermediate between the species. A geometric morphometric analysis shows the developmental trajectories from newborn to adult ringed (circles, *n* = 49) and gray (squares, *n* = 66) seals for the first three components (PCA). Arrows display developmental trajectories obtained by multivariate quadratic regressions of shape onto centroid size. The symbol size represents the relative centroid size of the crania and open symbols are male, black female. The larger size and more pronounced nasal region of gray seals are further developed in males. Arrowhead denotes the geometric mean calculated using the youngest ringed (*n* = 11) and gray seal (*n* = 11) crania. See Figure supp. 4, 5 and Materials and Methods.

Examining the details of cranial shape shows that the first two principal components distinguish features that separate the two species (explaining 61.6% of total variance) while the third component captures some mixed traits common to both of them (explaining 7.5% of total variance) (Figure supp. 5A). The hybrid differs from the geometric intermediate by having a shorter snout, narrower zygomatic arches and a more elongated braincase (Figure supp. 5B, C), agreeing with the original description reporting that some details of the hybrid cranial morphology are closer to one, or the other species (Lönnberg, 1929). We note, however, that the actual parents of the hybrid are not preserved, and thus some details of the morphology could represent one of the individual parents. Another factor affecting cranial shape is pronounced sexual dimorphism present in gray seals but not in ringed seals (Figure 4), although we did not detect dimorphism in newborn crania (*p* = 0.485 for gray seals and *p* = 0.839 for ringed seals, permutation tests on Procrustes distances with 10,000 rounds). Overall, the intermediate cranial and dental features of the seal hybrid analyzed suggest that a perfectly intermediate fossil specimen between two genera could potentially be a hybrid.

### Detecting hybridization in the wild

Finally, because the grey-ringed seal hybrid was born in captivity, in itself it does not imply that comparable hybridizations happen in the wild. However, as is the case for mammals in general (Polly et al. 2013; Shurtliff, 2013; Árnason et al., 2018; Warren et al., 2018), several seal species are well established as hybridising in the wild, including documentation of fertile intergeneric hybrids (Franco-Trecu et al., 2016), and a living intergeneric hybrid between the dentally disparate harp seal (*Pagophilus groenlandicus*) and hooded seal (*Cystophora cristata*) (Kovacs et al., 1997). Furthermore, behavioural observations have documented attempted matings by a wild gray seal male with harbour seal females (*Phoca vitulina*) (Wilson, 1975).

To estimate potential interbreeding in the history of wild gray and ringed seals, we examined introgression using the Patterson’s D-statistics approach (Green et al., 2010). In addition to Baltic gray seals and Baltic ringed seals, we sequenced genome wide data from Saimaa ringed seals (n = 12, the Weddell seal was the outgroup, see Materials and Methods). Saimaa ringed seals are a suitable contrast for the analyses (Figure 5A) because they have been isolated from the Baltic for approximately 9,500 years, with no documented presence of gray seals in Lake Saimaa (Valtonen et al., 2012; Ukkonen et al., 2014). The results show that Baltic ringed seals have a statistically significant excess of derived alleles shared with the gray seal compared to the Saimaa ringed seals (Figure 5B, Table supp. 10). We consider these results to be at least suggestive of interbreeding occurring in the Baltic between gray and ringed seals, and it is plausible that many seal populations and species will exhibit gene flow comparable to that in bears (Kumar et al., 2017) and horses (Jónsson et al., 2014). Taken together, the captive gray-ringed seal specimen can be considered to be a representative example of a phenotypically disparate hybridization.

**Figure 5.**
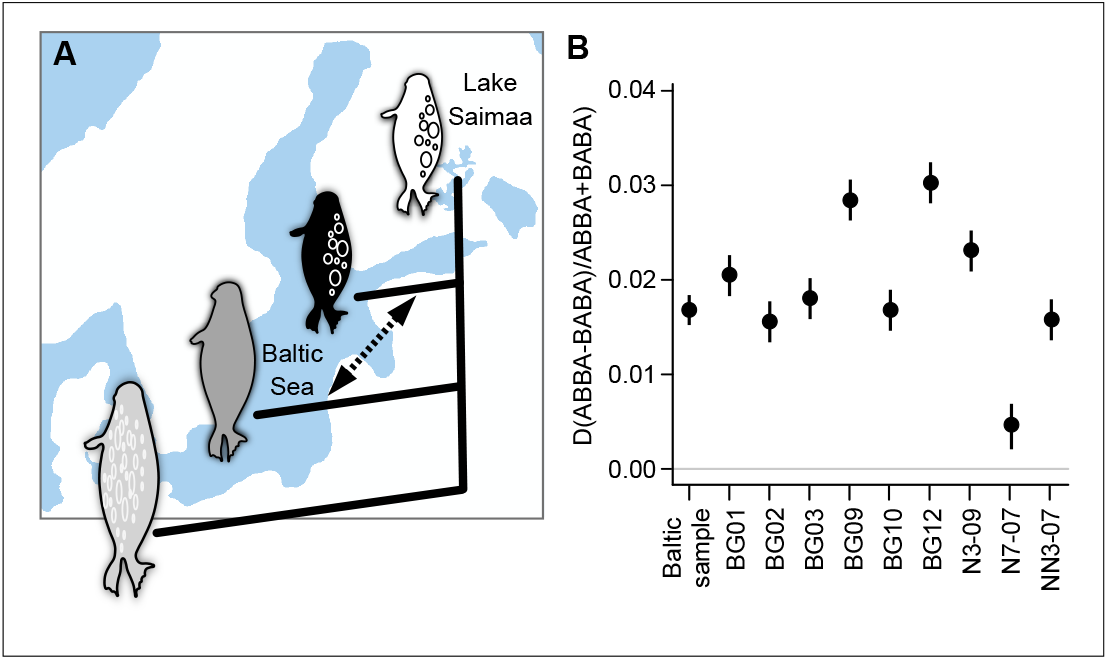
Introgression analysis suggests interbreeding between Baltic gray and ringed seals in the wild. **(A)** Saimaa ringed seals (white) have been landlocked in the Lake Saimaa, Finland for 9,500 years, providing a suitable comparison for Baltic ringed seals (black) and gray seals (gray). Outgroup was the Weddell seal (light gray). **(B)** Positive D-statistic values, calculated using D(Saimaa ringed seal, Baltic ringed seal; Baltic gray seal, Weddell seal) indicate gene flow between Baltic gray and ringed seals (arrow in A). Although the introgression and Z-scores vary between the individuals, all but one individual (N7-07) were *p* < 0.0001 (Table supp. 10 and Materials and Methods). Whiskers denote one standard error.

## Discussion

Gray and ringed seals have sufficiently morphologically different dentitions that, if they were to be discovered as unknown fossils, at least a genus level distinction could easily be justified. Yet, the combination of our genomic, phenomic, and developmental, analyzes show that these species can, and are likely to hybridize and that the resulting phenotype is a predictable combination of the two species. In more general terms, this hybrid bracket is indicative of the conservation of developmental mechanisms that tinker with quantitative traits. Whereas the overall conservation of developmental signaling in tooth development across the whole of jawed vertebrates is well established (Fraser et al., 2009; Jernvall and Thesleff, 2012), our results suggest interspecies and higher level conservation of the regulatory architecture underlying dental form. Previous reports have found increased occurrences of supernumerary teeth in mammalian hybrids (Goodwin, 1998; Ackermann et al., 2006), which could indicate that there is a point of divergence at which the developmental regulation begins to lose canalization. Loss of developmental regulation underlying extra teeth is supported by the frequent presence of supernumerary teeth in mice with null mutations affecting tooth development (Jernvall and Thesleff, 2012). Supernumerary teeth are relatively common in seal populations (Cruwys and Friday, 2006; Miller et al. 2007), but these are usually attributed to relaxed selection due to the lack of refined occlusion in seals (Cruwys and Friday, 2006; Miller et al. 2007; Jernvall and Thesleff, 2012). The potential role of interbreeding in the occurrence of extra teeth in seals remains to be determined.

In the case of paleontological research where phenotypic analyses are the basis for taxonomic inferences, our results suggest that many closely related genera could potentially hybridize, and that intermediates between distinct species may be hybrids themselves. The large morphological differences relative to the relatively modest genetic distance between gray and ringed seals are suggestive of an adaptive radiation phase of evolution, and we postulate that phenotypically disparate hybridizations are most likely to be observed in such radiations. In itself, hybridization between species has been also proposed to facilitate adaptive radiations (Seehausen, 2004; Warren et al., 2018). Finally, in our case, the hybrid seal was of the first generation, and continuing interbreeding would result in segregation of traits, something that might appear as a mosaic mixture of traits in the fossil record (Warren et al., 2018). With advancing understanding of the developmental basis of organ form, it may be possible to provide predictions of potential hybrids, even in cases where the phenotypic differences are substantial.

## Materials and Methods

### Specimen preparation

The hybrid was a newborn and although the crown morphogenesis of the permanent teeth was completed, the teeth were erupting and only partially mineralized. Seal deciduous dentition is vestigial. The right side dentition was removed from the jaw by Lönnberg (1929), and most of the teeth were cracked or split into halves during storage. The skull, jaws and isolated teeth were microCT scanned using a custom-built microtomography system (Nanotom 180 NF, phoenix|X-ray System and Service GmbH, General Electric, Wunstorf, Germany) at the Department of Physics, University of Helsinki, Finland. The voxel size was 32.1 μm for the skull and 16.6 μm for the tooth row. Volume data were processed in Avizo 9.0 (FEI Visualization Sciences Group) and surfaces were extracted and teeth segmented from the jaw using Artec Studio 9.0 (Artec 3D). Cracked tooth halves were manually reattached using Artec Studio 9.0. The tooth halves were joined together without deforming the meshes (Figure supp. 1). Skull images and drawings are from specimens in Finnish Museum of Natural History (Helsinki, Finland) and partly for *Homo* from (Tattersall and Schwartz, 1998).

### Seal genetic data

Hybrid DNA was isolated from the pulpa of a poscanine tooth that had fallen from the specimen in storage using NucleoMag kit (Macherey-Nagel GmbH & Co, Germany). The same method was used to isolate DNA from Baltic gray seal, Baltic ringed, and Saimaa ringed seal muscle tissue samples. The Baltic seal samples were obtained from the collections of the Saimaa Ringed Seal Genome Project, University of Helsinki, and Natural Resources Institute Finland, whereas the Saimaa samples originated from the collections of the University of Eastern Finland, which has a permission from Finnish environmental authorities for taking possession of and storing tissues of the Saimaa ringed seal (permission number: Dnro VARELY/3480/2016). All libraries and sequencing were performed at the DNA Sequencing and Genomics Laboratory, Institute of Biotechnology, University of Helsinki, Finland. The hybrid seal genome was sequenced with 2 separate runs of Illumina HiScanSQ platform and 11 separate runs of Illumina MiSeq platform to ca. 100X raw sequencing coverage (Table supp. 1). The base calls in the Illumina HiScanSQ platform were converted into text format (FASTA-format with individual base quality scores) using the CASAVA toolkit (bcl2fastq version 1.8.3) provided by the manufacturer. The Illumina MiSeq platform employed a primary analysis software called MiSeq Reporter post-run that produced the base calls in text format.

For all the Illumina reads, base call accuracy and read length filtering (adapter cutoff) was performed with cutadapt (Martin, 2011) using minimum accepted base call accuracy of 90% and minimum post-filtering read length of 75 bp. The Weddell seal (*Leptonychotes weddellii*) draft genome (Broad Institute) was used as a reference genome for the downstream genetic distances analysis. The statistics and downloads for this reference are available at: http://software.broadinstitute.org/allpaths-lg/blog/?p=647.

### Admixture and introgression analyses

Short read data for Hybrid, Baltic gray seal, Baltic ringed seal, and Saimaa ringed seal were mapped to the Weddell seal reference genome. Similarly, the reference Weddell seal individual’s short read data was mapped to its reference genome. The mapping was conducted using bwa mem (v. 0.7.15), samtools rmdup (v. 1.3.1) and GATK IndelRealigner (v. 3.7). The resulting bam files were used for introgression and admixture analyses.

To examine the hybrid, an admixture analysis was carried using NgsAdmix (Skotte et al., 2013) and the pipeline at: http://www.popgen.dk/software/index.php/NgsAdmix. To reduce computational burden, a random sample of 3.6 million markers (10% of the data) were used to reduce the computation burden. NgsAdmix was run 10 times with K=2 clusters on the hybrid, gray and ringed seals. Each of these runs converged to identical clustering.

Introgression was studied using Patterson’s D-statistics (Green et al., 2010) that, other than requiring the ancestral population to be randomly mating, is robust to variables such as variations in population size (Durand et al., 2011). D-statistics was calculated by abbababa2 module of ANGSD (Korneliussen et al., 2014) using the first 1,000 scaffolds of the Weddell genome (over 50% of the genome) and the default pipeline at: http://www.popgen.dk/angsd/index.php/Abbababa2. To examine individual differences, the D-statistics were calculated for each Baltic individual in addition to the Baltic sample. The Saimaa ringed seals were set as P1, Baltic ringed seals as P2, and Baltic gray seals as P3. The aligned sequences are available through the European Nucleotide Archive under accession number PRJEB25679.

### Genetic distances

To obtain a relatively robust and comparable measure of genetic distances between the hybridizing species pairs, we analyzed fourfold degenerate (ffd) sites from a large number of orthologous genes. As substitutions on ffd sites do not alter the encoded amino acid, they can be considered a proxy of the neutral evolutionary process and thus be relatively free of biases due to selection. A list of single-copy human genes and their orthologs in cat, dog, horse and chimpanzee were fetched from Ensembl BioMart (v. 83) (Kinsella et al., 2011). 4,826 complete gene sets with unambiguous orthology (“Homology Type” is “ortholog_one2one”; “Orthology confidence” is “high”) were retained and, for each reference species, nucleotide and peptide sequences of every transcript were fetched using the Ensembl REST API (Yates et al., 2014). Using last (v. 658) (Kiełbasa et al., 2011), scaffold-level genome assemblies of tiger, Weddell seal and donkey were aligned against the genomes of cat, dog and horse, respectively. Lift-over chains were built using the Kent source utilities (Kent et al., 2002). Short read data for donkey and lion were mapped to the horse and tiger genomes using bwa mem (v. 0.7.15), samtools rmdup (v. 1.3.1) and GATK IndelRealigner (v. 3.7) (Li and Durbin, 2009; McKenna et al., 2010). Locally generated short read data for Baltic gray seal and Baltic ringed seal were mapped to the Weddell seal genome. The data for the Neanderthal human individual were obtained as bam alignment files. Data sources are listed in Table supp. 2.

The gene annotations were divided into individual coding exons, removing split codons where necessary, and the exon coordinates in reference species (human, cat, dog, horse) were transferred to related non-reference genomes using CrossMap (v. 0.2.4) (Zhao et al., 2013). For each gene transcript, exons were independently extracted from the non-reference genomes using samtools (v. 1.3.1) and matched against the reference peptide sequences using Pagan’s (v. 0.61) (Löytynoja et al., 2012) translated pileup alignment. The resulting back-translated nucleotide alignments were flattened and degapped, combining all successfully extracted exons into one sequence. For species with bam alignment data (donkey, gray and ringed seal, Neanderthal), the genome regions surrounding the exons were reconstructed by placing observed sequence changes to the genome sequences of horse, Weddell seal and human. For each exon separately, variants were inferred with samtools mpileup and bcftools call (v. 1.3.1), and then placed to the genome sequences with vcfutils vcf2fq, requiring sequencing depth of 10 or higher. The exon regions of each transcript were extracted and combined into one sequence.

Sequence data were obtained for 4,329 genes. These sequences were aligned, selecting the longest transcripts, with Pagan’s translated pileup alignment. To remove artefacts of low quality sequence (for example, due to assembly errors), resulting multiple sequence alignments were filtered and regions where two neighboring codons contain more than one non-identical base among all the study species were masked. For different species sets, gene sequence alignments of at least 100 columns in length after filtering and removal of sites with missing data were concatenated, and ffd sites were extracted using R package rphast (v. 1.6.5) (Hubisz et al., 2010). Pairwise genetic distances were computed using R package ape (v. 4.0) (Paradis et al., 2004) and model T92, and phylogenetic trees (Figure 2B) inferred with RAxML (v. 8.2.9) (Stamatakis, 2006) using model GTRGAMMA. In addition, we measured the distances using all the codon third sites, and the relative distances remained largely the same.

The species set analyzed are: all twelve species, eight species known to hybridize, the two seals with each pair of hybridizing species. The numbers of input genes, codons and ffd sites as well as the pairwise genetic distances for different species sets are given in Table 1 and Tables supp. 3, 4.

### Dental material and analyses

Baltic gray and ringed seal material was used in comparisons to the hybrid as the parents were reported to be from the Baltic Sea (Dataset supp. 1, Table supp. 5). Postcanine teeth of seals are laterally compressed resembling early mammalian morphologies, and the lateral views capture most of the shape variation (Jernvall, 2000). Images were taken from the lingual views from a distance (approximately ten times the length of the tooth row) and the number of cusps and top-cusp angles were tabulated using ImageJ (version 1.51). The angle captures the relative height of the tallest cusp, and correlates with cusp number because cusp spacing and number are developmentally linked (Jernvall, 2000; Salazar-Ciudad and Jernvall, 2010). For teeth that had only one or two cusps, the line was drawn to the maximum anterior or distal ends at the crown base. We tabulated the presence of all cusps, including small incipient cusplets (arrowheads in Figure supp. 1B). All the measurements reported are from the right side. The pattern of results remains the same for the left side. Of the 65 gray and 65 ringed seal jaws studied, angle measurements were excluded from teeth with substantially cracked or worn cusp tips. We also excluded jaws with anomalies such as extra teeth. A sample of gray and ringed seal teeth were imaged twice and measured to confirm the robustness of the angle measurements (mean absolute error = 1.0 and 1.5, SD = 0.92 and 1.08, n = 40 and 46 teeth for gray and ringed seals, respectively). To test how intermediate the hybrid is between the species, we calculated average top-cusp angles between each gray-ringed seal pair, and tabulated the probability of obtaining the hybrid value from the gray-ringed seal averages (Figure 3B, Figure supp. 2). For visualization, dental specimens were scanned with PlanScan laser scanner (PlanMeca, Helsinki, Finland).

### Developmental modeling of teeth

We used ToothMaker (Harjunmaa et al., 2014) to simulate seal tooth development. The model integrates experimentally inferred genetic interactions with tissue biomechanics to simulate tooth development (Salazar-Ciudad and Jernvall, 2010), and here we used a version that had been experimentally explored in connection with mouse and vole molar development (Harjunmaa et al., 2014; Renvoisé et al., 2017). We focused on three parameters to simulate development of seal dentitions. To regulate spacing of cusps, the key difference between gray and ringed seals, we adjusted the strength of the enamel knot-secreted inhibition of enamel knot formation (parameter *Inh*). Experimental evidence on mouse molar development has shown how adjusting inhibition of new enamel knots results in quantitative changes in cusp spacing (Harjunmaa et al., 2014). Another strategy to alter cusp initiation would be to adjust the activator (Harjunmaa et al., 2012), but here we prefer the inhibitor due to the experimental data (Harjunmaa et al., 2012; 2014), and because increasing inhibition alone, which leads to larger spacing of cusps as in the gray seal, does not inhibit growth directly (only indirectly by inhibiting new enamel knots). The other two parameters used to simulate the species differences are epithelial growth (*Egr*) and anterior bias (*Abi*). Egr affects the relative growth of the epithelium and pointedness of cusps, and Abi the anterior extension of the tooth germ; both have been found previously to account for variation in species differences (Salazar-Ciudad and Jernvall, 2010; Harjunmaa et al., 2014; Renvoisé et al., 2017). Lateral biases (*Lbi* and *Bbi*) affect lateral expansion of the tooth and indirectly cusp position laterally (Renvoisé et al., 2017). Because seals have laterally compressed teeth, these parameters were kept constant. One candidate gene for the lack of lateral expansion in seals is a transcription factor Foxi3 (Jussila et al., 2015), as *Foxi3* haploinsufficient dogs have seal like lower molars (Kupczik et al., 2017). All the used parameter values are listed in the Table supp. 6. Top-cusp angle measurements were done using ToothMaker and manually (Figure supp. 3B). We modeled gray and ringed seal tooth rows by making constant parameter changes from tooth-to-tooth to mimic gradual shape changes along the jaw. All the simulations were run for the same number of iterations.

To simulate hybridization, for each tooth, we averaged the parameter values between the species, adjusted the averages 10% towards each of the parents, or used the gray or ringed seal value in different combinations (Figure supp. 3B, Table supp. 7). These different combinations basically simulate how additively the parameters function, but do not imply that each parameter would have one underlying locus. Rather, we consider the simulations to approximate the overall developmental effect of the genetic backgrounds. ToothMaker software, is freely available for Windows, MacOS, and Linux (available from the authors and with default seal parameters at https://github.com/jernvall-lab/ToothMaker).

### Analysis of the crania

31 ventral and 22 dorsal 3D landmark coordinates were acquired from skulls of complete ontogenetic series (from newborns to adults) of gray and ringed seals from the collections of Finnish Museum of Natural History (Helsinki, Finland) using a Microscribe G2X digitiser with 0.2 mm accuracy (Immersion). These data were merged together into a full configuration of 46 landmarks using FileConverter (Klingenberg, 2011) (Figure supp. 4). Digitizing was performed twice and one-way ANOVAs were used to test for digitizing error (Table supp. 9). We analysed variation among individuals (symmetric component of variation) apart from asymmetry (differences between left and right sides of individuals) (Savriama and Klingenberg, 2011). A Generalized Procrustes Analysis (GPA) was used to simultaneously superimpose all configurations and extract shape data by removing the effects of scale, orientation and position (Klingenberg, 2010). Centroid size, the measure of size in geometric morphometrics, was computed as the square root of the sum of the squared distances of all landmarks from their centroid (Dryden and Mardia, 2016). Permutation tests were used for testing differences in mean shapes using function ‘permudist’ from ‘Morpho’ R package (Schlager, 2017).

A Principal Component Analysis (PCA) performed on the covariance matrix of shape variables was used to show the main patterns of morphological variation between the two species. A 3D microCT-scan of the hybrid was warped to the consensus using the Thin Plate Spline (TPS) method via the function ‘warpRefMesh’ in Geomorph (Adams et al., 2017) and the main patterns of shape changes were visualised as deviations from this warped consensus for the first three principal components (PCs) (Figure supp. 5A). The hybrid was compared to the geometric morphometric mean and differences were visualised via warping and heatmaps (Figure supp. 5B and C). A multivariate quadratic regression of shape onto centroid size was used to investigate the main patterns of allometric growth for each species that were visualised with warped 3D microCT-scans of a newborn gray and ringed seal using Landmark Editor 3.6 and function ‘warpmovie3d’ (Figure 5, Movie supp. 1) (Wiley et al., 2005; Gerber and Hopkins, 2011; Schlager, 2017). All analyses were performed with MorphoJ 1.06d (Klingenberg, 2011), custom R (R Core Team, 2009) and Python scripts.

## Acknowledgements

We thank S. Mallick and S. Martin for discussions and advice on genome related analyses, J. Laakkonen, M. Kunnasranta, and M. Olsen on seal biology. We thank the members of the Saimaa Ringed Seal Genome Project for help, for collections access and specimen loans we thank I. K. Hanski and E.-S. Hyytiäinen from the Finnish Museum of Natural History (Helsinki, Finland) and D. Kalthoff from the Museum of Natural History (Stockholm, Sweden), who also helped us to locate the hybrid specimen. We thank UH librarians for help with old reprints, and H. Suhonen, A. Kallonen, M. Christensen, and J. Muszynski for tomography and 3D data processing. We also thank P. Timonen and S. Oksanen for help with Baltic seal samples. We thank CSC-IT Center for Science, Espoo, Finland, for computing facilities. The personnel at DNA Sequencing and Genomics Laboratory, Institute of Biotechnology, University of Helsinki is acknowledged for excellent technical work. We thank PlanMeca corporation for equipment loan. We also thank Natural Resources Finland for providing samples and Metsähallitus Parks & Wildlife Finland for co-operation. Key financial support was provided by Jane and Aatos Erkko foundation and the Academy of Finland.

## Author Contributions

J.J. and P.A. conceived the project, I.J.C. identified and Y.S. reconstructed the hybrid specimen, P.A. sampled DNA from the hybrid, M.V. collected and M.V. and J.J. analyzed dental data, T.J.H., I.S.-C. and J.J. performed the developmental simulation, Y.S. collected and analysed cranial data, S.G. performed the allometric analyses for cranial data, J.K, O.-P.S., A. Lyyski, L.P., and P.A. performed the sequencing and J.K., O.-P.S, and L.H. the processing of the reads, A.L. analyzed genetic distances, and P.R. and A.L. analyzed introgression. J.J. wrote the initial manuscript and all authors discussed the results and provided input and scientific interpretations on the manuscript.

## Data and materials availability

The aligned sequences are available through the European Nucleotide Archive under accession number PRJEBxxxxx with individual sample accessions (xxx…xxx). The cranial specimens listed in this study are archived in the Finnish Museum of Natural History and Swedish Museum of Natural History. Any further data not available in the Supporting Materials are available from the corresponding authors on request.

